# LncPlankton V1.0: a comprehensive collection of plankton long non-coding RNAs

**DOI:** 10.1101/2023.11.03.565479

**Authors:** Ahmed Debit, Pierre Vincens, Chris Bowler, Helena Cruz de Carvalho

**Affiliations:** Institut de Biologie de l’ENS (IBENS), Département de biologie, École normale supérieure, CNRS, INSERM, Université PSL, 75005 Paris, France; Université Paris Est-Créteil (UPEC), Faculté des Sciences et Technologie, 61, avenue du Général De Gaulle, 94000 Créteil, France

## Abstract

Long considered as transcriptional noise, long non-coding RNAs (lncRNAs) are emerging as central, regulatory molecules in a multitude of eukaryotic species, from plants to animals to fungi. Yet, our knowledge about the occurrence of these molecules in the marine environment, namely in planktonic protists, is still elusive. To fill this gap of knowledge we developed LncPlankton v1.0, which is the first comprehensive database of marine plankton lncRNAs. By integrating the predictions derived from ten distinctive coding potential prediction tools in a majority voting setting, we identified 2,210,359 lncRNAs distributed across 414 marine plankton species from over nine different phyla. A user-friendly, open-access web interface for the exploration of the database was implemented (https://www.lncplankton.bio.ens.psl.eu/). We believe LncPlankton v1.0 will serve as a rich resource for studies of lncRNAs that will contribute to small- and large-scale analyses in a wide range of marine plankton species and allow comparative analysis well beyond the marine environment.

## INTRODUCTION

Often referred to as the “dark matter” of genomes, the non-protein coding DNA portion of eukaryotic genomes has, in the last decade, been shown to be much bigger than what could have been anticipated. Furthermore, with the advent of deep sequencing it has become clear that genomes are pervasively transcribed and that the non-coding fraction generates huge amounts of transcripts that cannot be accounted for as simple “junk” or transcriptional noise (1). Among this non-coding fraction, long noncoding RNAs (lncRNAs) represent the most abundant and prevalent class (2, 3). LncRNAs are arbitrarily defined as transcripts of more than 200 nucleotides (nt) in length that lack consensual open reading frames (2). Like mRNAs, lncRNAs are largely polyadenylated, capped and processed (2, 3). However, besides their lack of protein coding potential, lncRNAs exhibit certain characteristics that distinguish them from mRNAs. These include a low GC content, low number of exons, short sequence length, low sequence conservation, and low expression levels (4). In terms of function, lncRNAs have been shown to play important regulatory roles in various biological processes and diseases, namely cancers, which involve the control of epigenetic modifications as well as gene and protein expression regulation (1, 3).

With the booming of high-throughput sequencing techniques and the exponential rise of transcriptomics data in public repositories, several databases dedicated to lncRNAs have been developed. Some are exclusively devoted to human lncRNAs, like GeneCaRNA (5), and LNCipedia (6), while other databases have grouped together a significant number of lncRNAs coming from photosynthetic organisms such as CANTATAdb (7) and GREENC v.2 (8). The NONCODE knowledge database v6.0 on the other hand compiled lncRNAs from 39 species including 16 animals and 23 plants (9) and the RNAcentral catalogue is an extensive database of lncRNA sequences from a broad range of organisms (10). Despite such extensive work on the identification of lncRNAs in animals and plants, the ocean remains largely unexplored. To fill this gap of knowledge, our study took advantage of the publicly available transcriptomic data originating from more than 400 marine protists from the Marine Microbial Eukaryote Transcriptome Sequencing Project MMETSP (11), to perform a thorough screening to identify and annotate lncRNAs in marine plankton. In order to do this, a computational pipeline using a majority voting-based ensemble learning technique was developed and applied. The objective was to increase the reliability and promote the diversity of the ensemble model. Furthermore, a joint prediction is likely to behave better than any single model as recommended by Duan Y. et al. (12), and likely obtain higher cross-species prediction performance (13). In order to use the strength of multiple classifiers and to enhance their performance, several ensemble methods have been developed in recent years, such as TLClnc (13), which combined a stacking of SVM predictors and a naïve Bayes classifier. Conversely, Simopoulos et al. (14) proposed a prediction method based on the stochastic gradient boosting of random forest classifiers, and LncRNApred (15) proposes a method using hybrid features. To promote the reliability of the results, CRlncRC (16) was developed on the basis of five machine learning models including Random Forest (RF), Naïve Bayes (NB), Support Vector Machine (SVM), Logistic Regression (LR) and K-Nearest Neighbors (KNN), to predict cancer-related lncRNAs. In our study, we combined ten coding potential prediction classifiers (see Supplementary Table S1) into a single meta-learner. The selected tools used different AI algorithms, provided models pre-trained with diverse fine-tuned features, were non-species specific and achieved good performance within a reasonable runtime. Furthermore, two tools considering full and partial sequence length were included, mRNN (17) and LncADeep (18). In addition, the tools selected presented a good usability score, which is based on ease-of-use in installation and running of the tool as described in (19). Briefly, each transcript was predicted by each classifier and labelled as “Coding” or “Noncoding”, and the class with the highest number of votes was the outcome. Several filters were applied on the “Noncoding” set to discriminate between lncRNAs and sncRNAs.

The predicted lncRNAs and their annotations, including the nucleotide sequence, the encoded ORF(s), the folding energy, and the predicted secondary structure were organized and stored in a database, LncPlankton, with the aim to update it periodically on the basis of new knowledge and a potential expansion in the number of planktonic species screened. A user-friendly, open-access web interface for the exploration of the database was also implemented (https://www.lncplankton.bio.ens.psl.eu/). The user interface provides modules for browsing, searching, and downloading lncRNA data per species and/or per phylum, as well as interactive graphs, and an online BLAST service. Additionally, a SHINY application was integrated which allows the user to customize and visualize the classifications. With this user-friendly interface, we anticipate that LncPlankton will provide a rich source of information about lncRNAs for small-and large-scale studies in a variety of plankton taxa and contribute significantly to future efforts aimed at deciphering the biology and evolution of lncRNAs in diverse eukaryotic lineages.

## MATERIAL AND METHODS

### Data sources

In the current version of LncPlankton (version 1.0), transcriptomic data of 406 marine micro-planktonic species derived from the Marine Microbial Eukaryote Transcriptome Sequencing Project (MMETSP) (11) were used. The most represented phylum is Bacillariophyta (diatoms), to which we have additionally added the transcriptomes of six other cosmopolitan diatom genera, originally not included in the MMETSP project. Furthermore, the transcriptomes of the two reference diatom species, *Phaeodactylum tricornutum* (20) and *Thalasiossira pseudonana* (21), were also included and assembled using an in-house assembly pipeline (Figure S1). Taken together the transcriptomes screened covered > 9 phyla (Figure 1A). In total, 11,623,179 contigs were obtained across 414 species, almost half of which belonging to the Dinophyta and Bacillariophyta groups, with 3.4 and 2.4 million contigs, respectively (Figure 1A). The average contig length varied between 593nt (Dinophyta) and 949nt (Cercozoa) with a median = 651nt across all species (Figure 1B).

**Figure 1.**
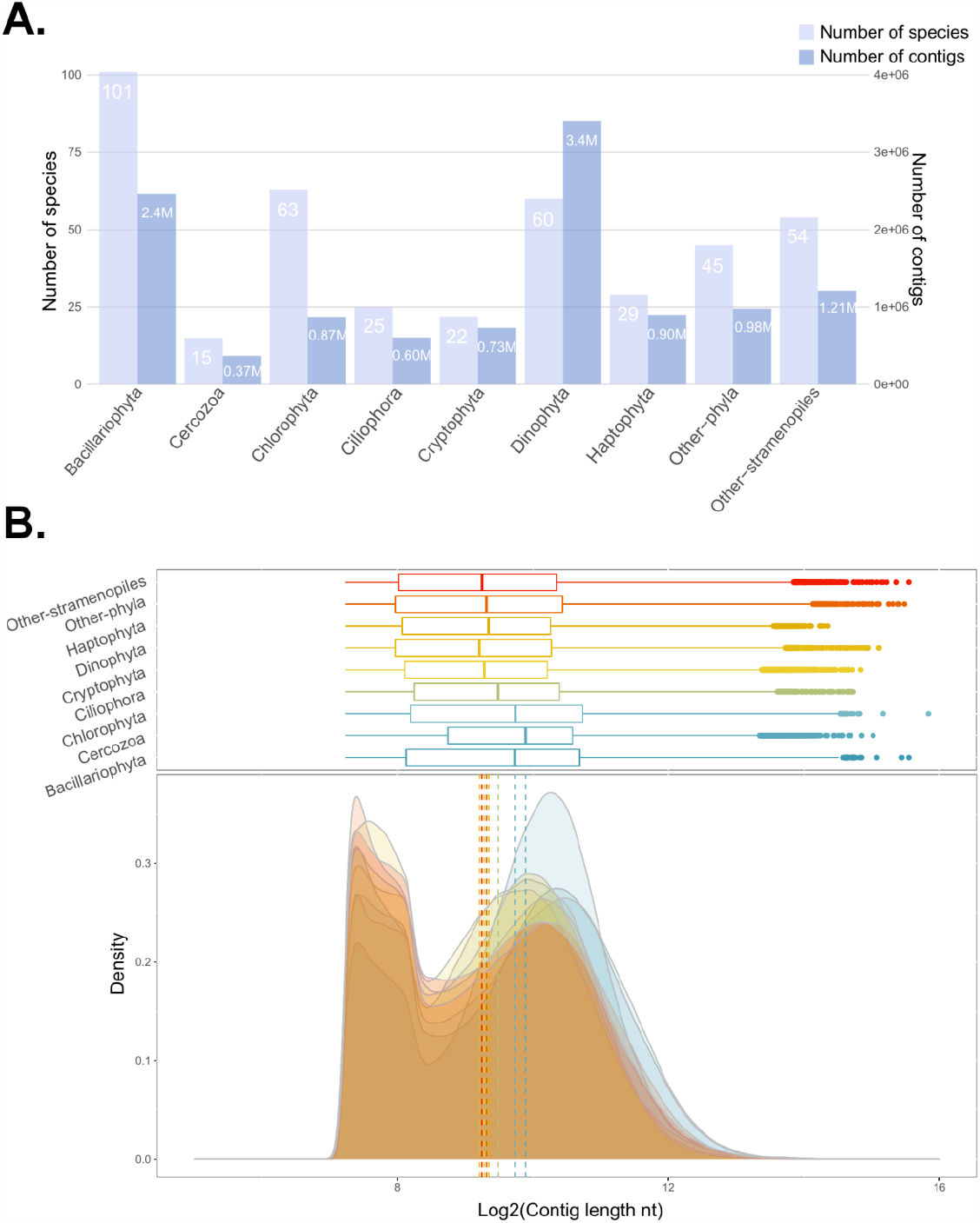
Data used in LncPlankton v1.0. **(A)** The distribution of the number of species and the number of assembled contigs across the different phyla; the number of contigs is displayed in millions and reported on the right scale y-axis of the graph and **(B)** The density distribution of the scaled log2 contig length across phyla, the dotted vertical lines denotes the median length.

### Data analysis pipeline

The assembled transcriptomes (in a *FASTA* format) provided by the MMETSP project (11) were screened for the prediction and identification of lncRNA-like transcripts, in each species, using the majority voting-based procedure inspired by the ensemble machine learning methods. The aim was to increase the reliability and promote the diversity of the ensemble model by combining all the predictions from multiple models. To achieve that, 10 protein coding potential prediction tools were selected and used in the pipeline. Each tool used a different algorithm and different training features, and may produce a different classification result. Aggregating those different techniques in a majority setting led to a single prediction representing the group“s consensus. We believe that such a hybrid technique leads to more reliable results and acceptable performance.

We selected the 10 tools according to the following criteria: i) the tools must use a different AI approach: among the 10 tools, 3 used a SVM-based model CPC2 (22), LncFinder (23), and longdist (24), 2 were based on XGBOOST LncDC (25) and RNAmining (26); 2 on maximum likelihood estimation MLE, and logistic regression LGC (27) and CPAT (28), respectively; and 3 other tools were based on deep learning approaches with different intrinsic architectures; convolutional neural network CNN in RNASamba (29), Recurrent neural network RNN in mRNN (17), and deep belief network DBN in LncADeep (18); ii) the tools must provide pre-trained models with diverse fine-tuned classification features; and iii) the tools should achieve good performance, particularly for cross-species prediction within a reasonable ease-to-use score according to the usability score defined in (19). In addition, the tools should accept sequences presented in a *FASTA* format. The tools selected and the key characteristics of each of them are summarized in supplementary Table S1.

The selected tools were applied using their default parameters and their default pre-trained classification models. The Python package *ezLncPred* (V.1.0) (30) was used to run the following tools: CPC2, CPAT (*-p Human*), LGC and longdist. The remaining tools were run using an in-house script with the following parameters: LncADeep (*-MODE lncRNA*, and the coding potential probability of a transcript was calculated as the mean of probabilities generated by the 21 intrinsic models of the algorithm), lncFinder (*svm*.*model=“human”*), mRNN (mRNN_ensemble which used the weighted average prediction of the five best single mRNN models), RNAmining (*-organism_name Homo_sapiens*), and RNASamba (Full and partial weighted models were included). The *FASTA* files containing the sequence contigs were used as an input to the pipeline displayed in (Figure 2). Basically, a transcript sequence was inputted to each tool which predicted both the forward and the reverse strand, and labelled them as “Coding-like” or “NonCoding-like”. A transcript was considered as “NonCoding-like” within a single tool only if both strands were labelled “NonCoding-like”, otherwise it was assigned the “Coding-like” label. For the assembly of the two species *Phaeodactylum tricornutum* and *Thalasiossira pseudonana*, stranded libraries were used, thus only the forward strand was assessed. The majority label output from all the tools was considered as the final coding potential class. Alongside the majority class, a non-coding potential score was calculated as the number of tools labelling the transcript as “NonCoding-like” divided by 10. This score was used to calculate the reliability (confidence level) of the lncRNA transcripts identified. The transcripts with significant hits in either the Pfam (V.35.0) or SwissProt (V.2023.01) databases were filtered out. A series of stringent filters was then applied to the “NonCoding-like” transcript sequences to classify them as long noncoding RNAs “lncRNAs” or small noncoding RNAs “sncRNAs” on the basis of transcript length and peptide length information.

**Figure 2.**
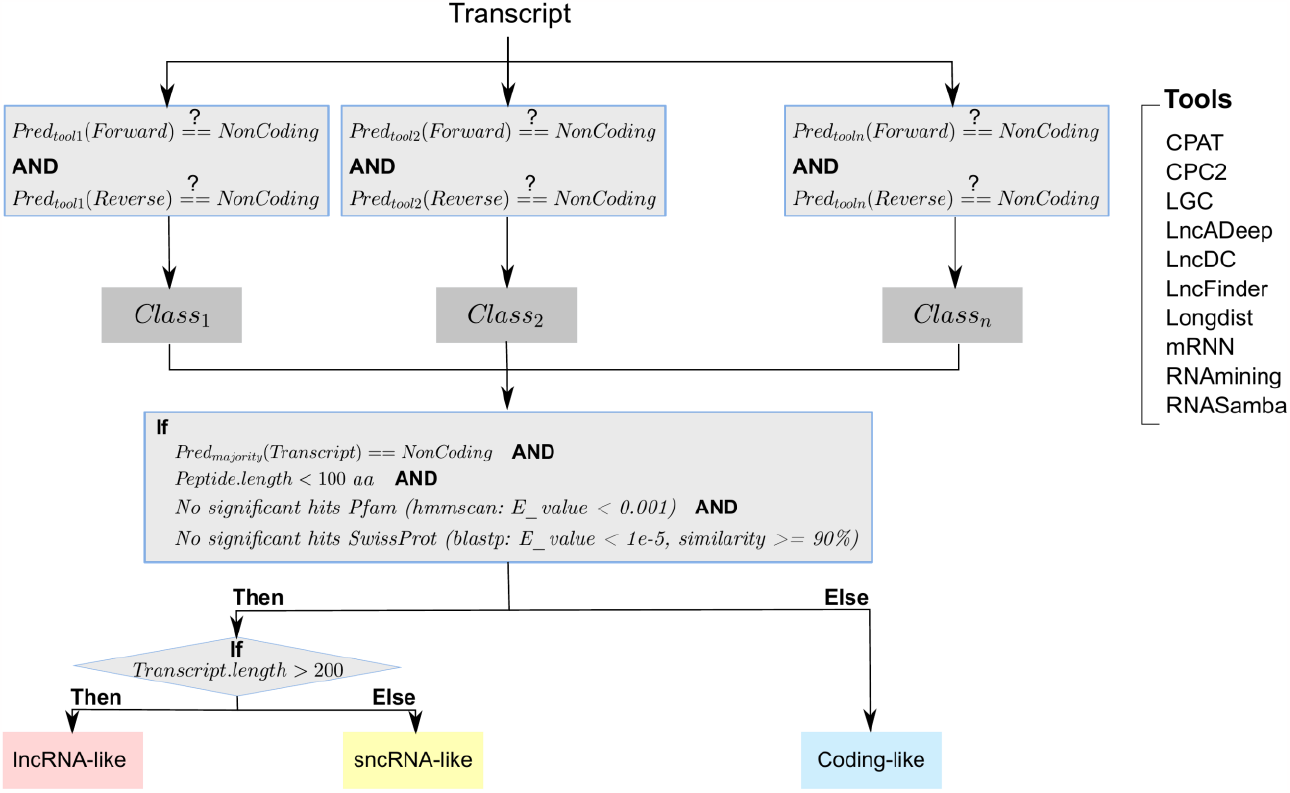
Overview of the majority voting-based pipeline for the prediction of lncRNAs. The 10 tools included in the procedure are displayed on the right panel.

### Evaluation of the performance of the majority voting-based procedure

The majority voting-based pipeline was tested on different sets of coding and non-coding datasets related to 18 different organism species and compared to the state-of-the-art coding potential prediction tools. Each tested dataset was independent of the training sets used in the construction of the pre-trained classification models related to each method. Among the 18 organisms tested, 10 datasets were perfectly balanced, containing the same number of coding and non-coding RNA transcripts. Furthermore, in order to test the robustness of the tools towards biased datasets, 7 highly imbalanced testing sets were considered. The description of the tested datasets is provided in the supplementary data (Table S2).

Each tool was applied to each tested dataset to predict the classes of the transcripts. The default parameters and pre-trained models were used for each tool. Tool outputs were parsed and analysed with custom R scripts. A cross-tabulation of observed and predicted classes was generated, and the performance metrics including the accuracy, sensitivity, and specificity were calculated using the *“confusionMatrix”* method of the caret package (https://CRAN.R-project.org/package=caret). The Receiver operating characteristic (ROC) curve as well as its AUC (Area under the curve) value was also computed using the ROCR package (http://rocr.bioinf.mpi-sb.mpg.de). All tested tools considered coding transcripts as positive and non-coding transcripts as negative sets.

### Architecture of the database

The 3-tier client/server architecture model containing data, logic, and presentation layers has been implemented for LncPlankton as shown in (Figure 3A). The data layer represents the data storage part which is handled by a relational database (Figure 3B) setup with the popular MySQL (version 5.7.36) open-source relational database management system (RDBMS). The data layer is expanded with NoSQL file storage. The logic layer represents the core of the architecture, and is responsible for the communication between the user queries from the presentation layer, fetching the data from the data layer, processing the data, and formatting the response to the presentation layer. The JSON-based (JavaScript Object Notation) data structure is mainly the most used format. In addition, the logic layer is integrated with the followings components:

**Figure 3.**
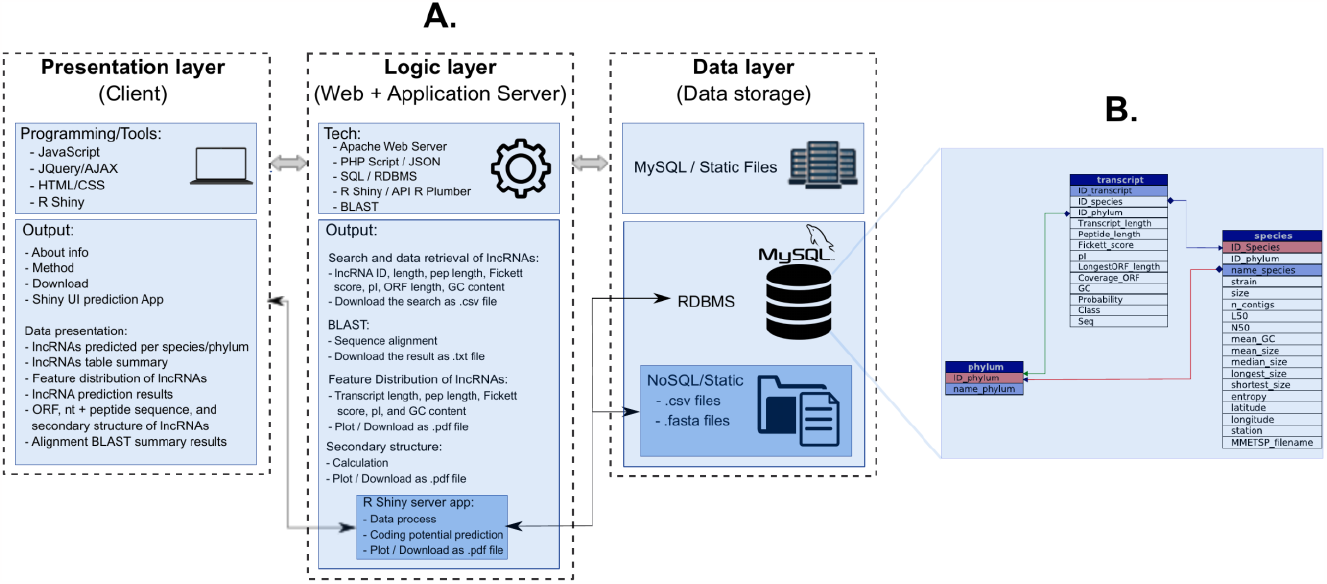
Schema of the Web/Database system of LncPlankton. **(A)** A detailed 3-tier architecture implemented in LncPankton; **(B)** The mySQL relational database scheme of LncPlankton containing three main tables: transcript, species, and phylum

- A BLAST program implemented via the interface rBLAST (https://github.com/mhahsler/rBLAST) for online similarity search,
- A RNAfold program implemented via the package LncFinder (23) and RNAPlot implemented via the package RRNA (31) for the calculation and the visualization of secondary structure,
- ORFfinder implemented via LncFinder (23) for the exploration of lncRNA containing sORFs, and seqinr package (https://github.com/cran/seqinr) for the translation to peptide sequences.

Those functionalities were provided via an express REST API web service implemented using the R package Plumber (https://github.com/rstudio/plumber). A Shiny server function was also developed and was integrated into the logic layer. This function processes the request of the shiny prediction app from the presentation layer and uses the static part of the data layer in addition to the SQL part. The presentation layer contains several modules based on AJAX (Asynchronous JavaScript and XML), jQuery (JavaScript Query system version 3.5.1), and the PHP server-side scripting language (version 7.1.26), as well as the CSS (Cascading Style Sheets) code to describe how HTML elements are to be displayed on user side web interface. JQuery and AJAX provide methods to perform asynchronous call requests to the logic tier using GET and POST methods, parsing the JSON response, and dynamically rendering the browser display.

The Web server is hosted on an Ubuntu (22.04.3 LTS) operating system using an Apache (version 2.4.52) web server. The user interface was tested and is functional across major web browsers including Chrome, Safari, and Firefox on Linux, Mac, Android, and Windows platforms. All graphs are generated dynamically using the open-source Chart.js library (version 3.5), and plotly R library (version 4.10.0).

### Implementation of the prediction application

The shiny application was built in R (V.4.3.1) using the shiny framework. The app currently depends on the following R packages: shiny, shinyWidgets, reshape2, wesanderson, dplyr, ggplot2, ggthemes, tidyverse, ggrepel, shinybusy, shinyjs, DT, plotly, leaflet, and RMySQL.

## RESULTS

### Performance of the majority voting algorithm

We compared the performance of the proposed majority voting-based method with the ten tools selected individually (Supplementary Table S1). We used as input all coding and non-coding RNA sequences from the testing datasets of the 18 species described in Supplementary Table S2. Our method showed the highest mean accuracy across all the datasets with a mean = 0.95 (Figure 4A), and low inter-dataset variability (variance: 0.0014). At the species level, our method outperforms the other tools in seven organisms, and performs well in cases where the other tools presented poor performances; for example, for *Homo sapiens* the majority voting procedure yields an accuracy of 0.95 while longdist and RNAmining yield only 0.56 and 0.54, respectively. For the remaining organisms, our method gets approximately the same accuracy as the other tools (Supplementary Table S3).

**Figure 4.**
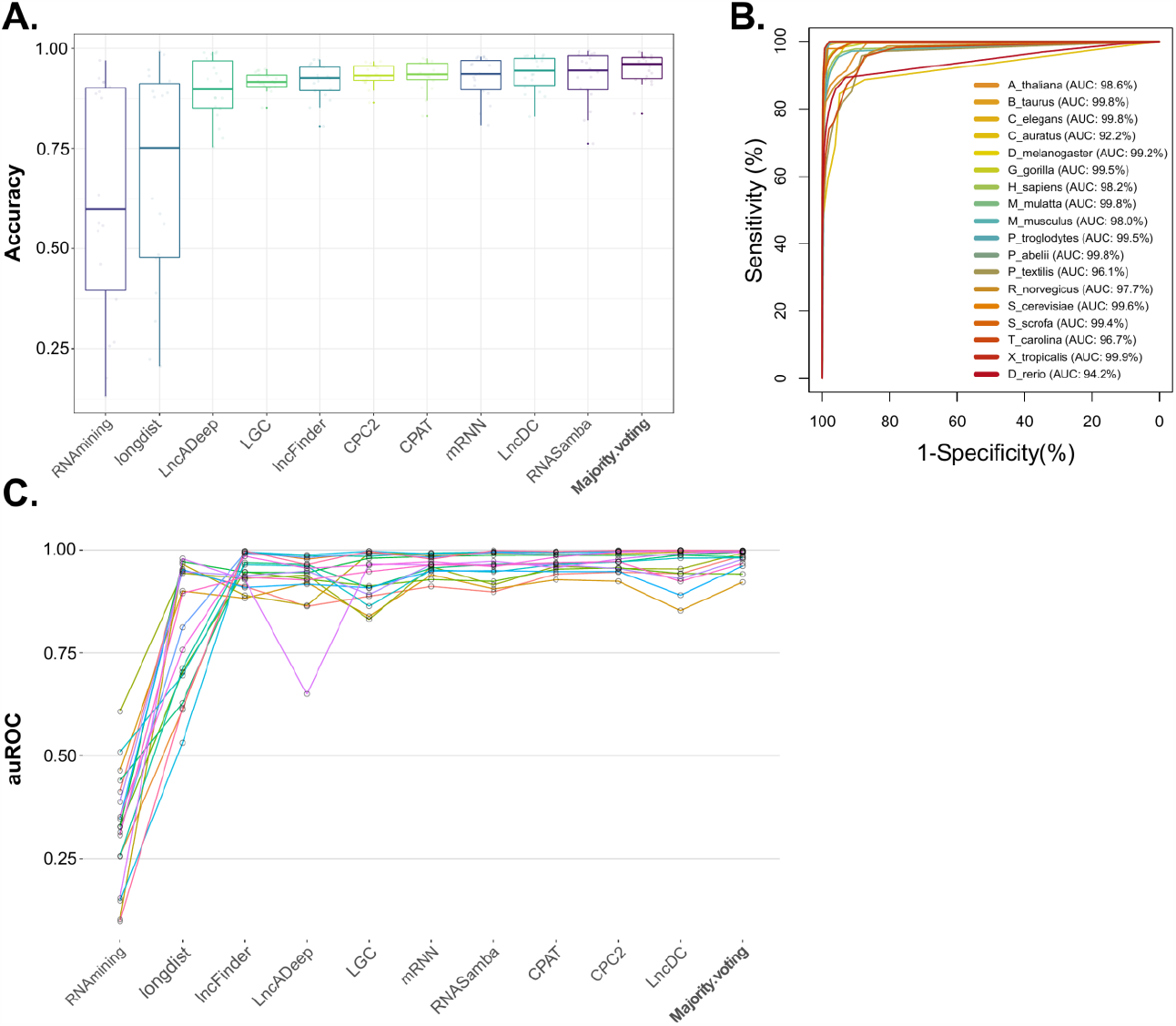
Benchmarking of ten coding potential prediction methods and the majority voting-based procedure using 18 independent testing sets. **(A)** Distribution of the accuracy across the 11 methods tested; the majority voting method on the last column showed the highest mean accuracy with a small variability; **(B)** ROC curves of the majority voting-based procedure on the 18 datasets tested, the AUC values corresponding to the curves are also reported; and **(C)** Distribution of the AUC scores obtained by each method on the 18 datasets.

In addition, our method showed comparable AUC performance (AUC: 0.92-1.00) with the best performing coding potential methods like CPAT and CPC2 (Figure 4C). Among the 18 datasets tested, our method showed the highest AUC score in 13 datasets (Supplementary Table 6). The AUC values obtained by our method for the 18 organisms are shown in Figure 4B.

The detailed results regarding the accuracy, the sensitivity, the specificity, and the AUC of all the tools can be found in Supplementary Tables S3, S4, S5, and S6, respectively.

### Data content of LncPlankton V1.0

Data in the current version of LncPlankton v1.0 was based on transcriptomic datasets from 414 planktonic species ranging from the Cercozoa, representing 3.6% of the total number of species screened, to the most abundant phylum Bacillariophyta, with 24.4% (Figure 1A). Using a majority voting-based method combining the 10 coding potential tools with strict prediction criteria (Figure 2), 2,210,359 lncRNA transcripts were identified from an input set of 11,623,179 transcripts. Among them, 239,116 were predicted as non-coding by all the tools and have a non-coding potential score = 1, we dubbed them “high-confidence lncRNAs” as their non-coding potential status was indisputable (Figure 5A). The distribution of the predicted lncRNA transcripts across the groups is shown in Figure 5B. More than 45% of the total lncRNA transcripts identified belong to the 2 most abundant groups, the Dinophyta (531,029 lncRNAs, 24.1%) and the Bacillariophyta (482,473 lncRNAs, 21.8%). At the species level, the highest percentage of lncRNAs were identified in the dinoflagellate *Karenia brevis Wilson* (23,686 lncRNAs, 1.1%) followed by the diatom *Fragilariopsis kerguelensis* (21,011 lncRNAs, 1%), while the lowest in the Rhizarian *Minchinia chitonis* (83 lncRNAs, 0.003%) and the dinoflagellate *Thoracosphaera heimii* (with only 10 lncRNAs) (see Supplementary Table S7).

**Figure 5.**
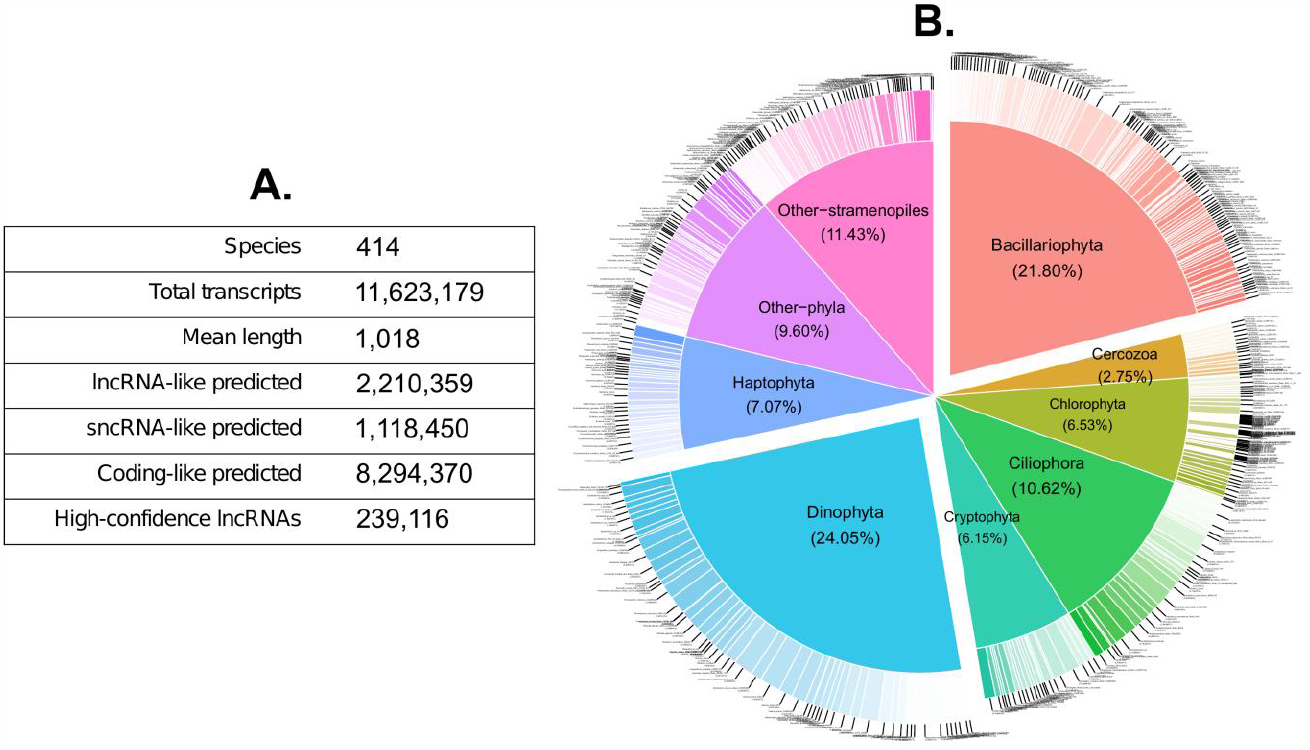
The current content of LncPlankton v1.0. **(A)** Summary statistics of the content of the database, and **(B)** The percentage of transcripts predicted as lncRNA-like by the majority voting pipeline distributed across phyla.

### Functional modules of LncPlankton V1.0

LncPlankton v1.0 provides a very useful user interface (UI) accessible from the browser. The UI offers various ways to browse and search lncRNA resources (Figure 6). Furthermore, users can download the data deposited in LncPlankton v1.0 in FTP bulk or programmatically through dedicated APIs.

**Figure 6.**
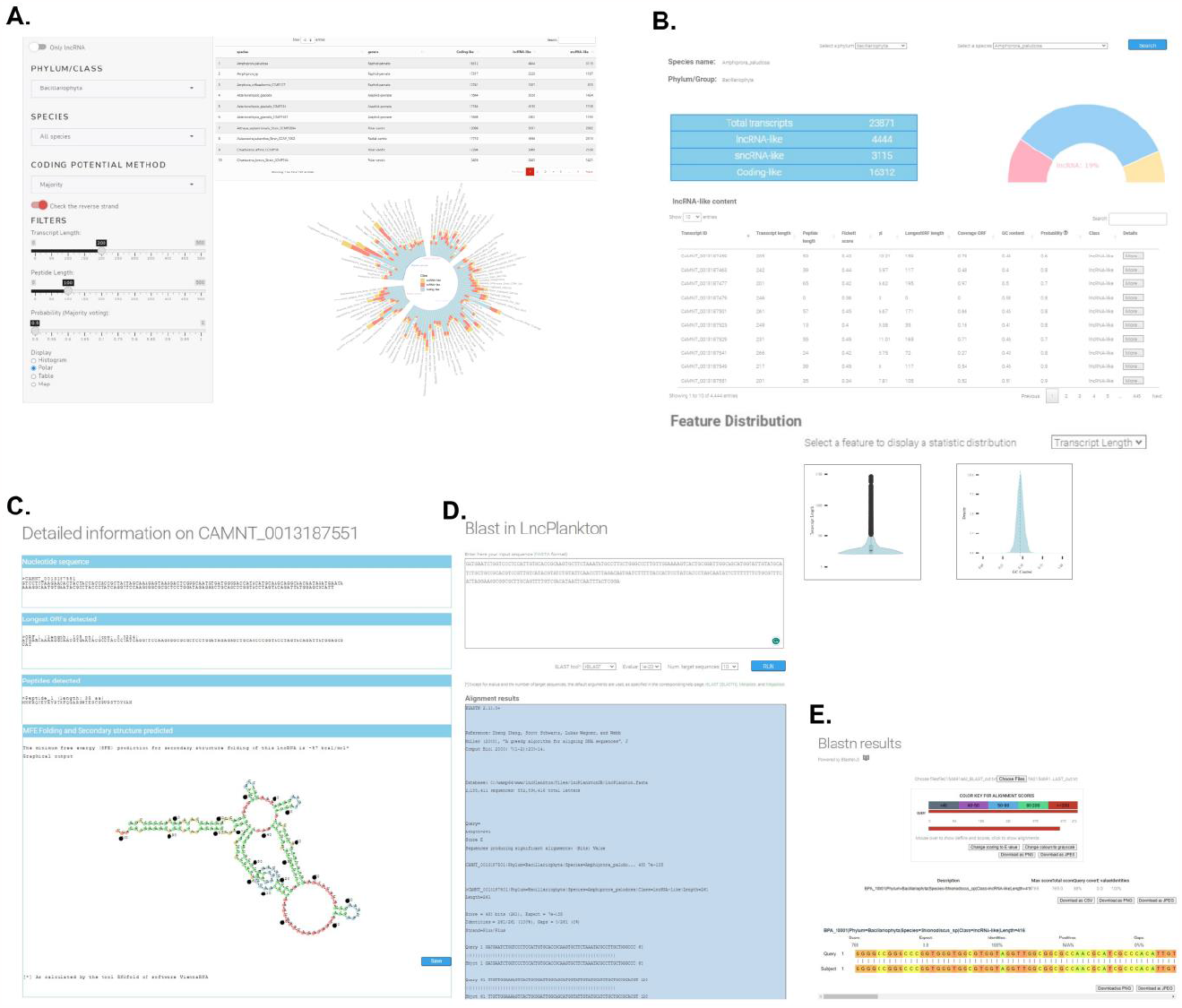
Screenshots of LncPlankton web interface and modules. **(A)** Interface of the LncPlankton prediction application. **(B)** Results page showing the content of the transcriptome of the diatom species *Amphiprora paludosa*. A table showing the features of all the lncRNAs predicted by the majority voting procedure was also displayed. **(C)** A page showing the detailed information on a predicted lncRNA including the secondary structure of the transcript **(D)** Online BLAST module and visual output in the LncPlankton database. Users can run blast against plankton lncRNAs by submitting the sequence in *FASTA* format, and then the result is displayed in text format. **(E)** The alignment result can be visualized in a more convenient way thanks to the BlasterJS module.

#### Search per species

The current release of LncPlankton allows querying a given species to explore the content of its transcriptome. The user is required to select a phylum and a species from drop-down lists. A page will then appear (Figure 6B) displaying a summary table with the number of transcripts classified as lncRNA-like, sncRNA-like, and Coding-like by the majority voting procedure described above. A pie chart reporting the percentage of lncRNAs found is also displayed. The user can also visualize the distribution of lncRNA features such as transcript length, peptide length, GC content and Fickett score. Additionally, all the information about the lncRNAs found for the selected species are reported in a table with one row per lncRNA. The presented data include lncRNA id, length, peptide length (if any), Fickett score, iso-electric point, longest ORF length (if any), coverage ORF (if any), GC content, class determined by the pipeline above, and the probability of coding potential generated by the majority voting procedure. To get access to more information on the selected lncRNA the user can click upon the “*More*…” button in the last column of the table.

#### lncRNA details webpage

The detailed information about the lncRNA selected is displayed on this page (Figure 6C). In addition to the basic details of lncRNA sequence (including phylum/group, species name, length, GC content, the majority voting probability, and the confidence level), the page displays the ORFs and conceptual translation products of the sequence, alongside with the length of ORFs detected, their coverage information, and the length of the translated peptides. Furthermore, the page shows the one-dimensional dot-bracket notation and the two-dimensional rendering secondary structure of the lncRNA. The Minimum Free Energy (MFE) of the structure folding was also calculated by the RNAfold tool (V.2.5.1) and displayed. A “save” button was added to allow the user to download and save the structure.

#### lncRNA prediction app

The lncRNA content of the current release of LncPlankton was generated using the majority voting-based procedure described above. To allow the user to customize their predictions, for example using a preferred coding potential prediction tool, and/or to modify the cut-off value of the filters, a SHINY application was developed and embedded to the UI. The application contains two panels (Figure 6A): i-The input panel in which the user can select the phylum, the species, and the coding potential tool from dedicated drop-down lists. Other inputs can be set by the user such as the transcript length, the peptide length, and whether or not to include the reverse strand in the prediction. The input panel offers also the possibility to select the manner in which the result will be displayed, namely as a histogram, polar plot, table, or map (with the sampling location coordinates of the selected species); ii-The output panel, which renders the results and displays it according to the choice of the user. All the figures and tables generated can be optionally downloaded and saved. The output panel was implemented using the plotly package (V4.10.1).

#### BLAST module

The user can perform a sequence-based search of data stored in LncPlankton using BLAST (Figure 6D). The input sequence should be in a *FASTA* format; and the user can select between two tools, BLASTN and MEGABLAST. Except for the expectation value (E-value) and the number of target hit sequences which can be selected in dedicated drop-down lists, the default arguments were used. The BLAST search outputs a raw report which includes pairwise alignment, BLAST hits based upon alignment scores and other measures of statistical significance. To interactively display the BLAST result, a viewer module was implemented using the BlasterJS library (32), and was integrated to LncPlankton (Figure 6E).

#### Download

Both the full and the high-confidence collections of lncPlankton can be found on the download page. The lncRNA collections related to each species can also be downloaded in a *FASTA* format separately. In addition, the majority voting R package used for the prediction can be retrieved from this page.

#### API

A representational state transfer application programming interface (REST-API) was implemented using the plumber package, and made available to allow programmers to interface with LncPlankton programmatically. The API returns documents in the JSON format and can be used in any programming language. In the current version of LncPlankton, three APIs were implemented and documented to access and use data:

- *GET /lncRNA_details:* information about lncRNA transcript including: sequence length, peptide length, reliability (confidence calculated by the majority voting-based procedure), GC content, ORF coverage, MFE, Fickett score, isoelectric point, nucleotide sequence, peptide sequence.
- *GET /search_per_species:* detailed information about species including the number and the percentage of transcript predicted.
- *GET /search_per_phylum*: detailed information about phylum including number of species, and statistics about the prediction of the transcriptomes of the species.

## DISCUSSION AND FUTURE DIRECTIONS

Determining the coding potential of a transcript is a crucial step in the identification of lncRNAs, yet it represents a complex task due to overlapping characteristics and functions that exist between coding and noncoding RNAs (33). To overcome this challenge, many computational methods have been developed. The majority of tools use features derived from the nucleotide sequence of the transcript such as the Fickett score, GC content, and kmer composition. Some of the tools like lncFinder use secondary structure properties to try to improve the discrimination between non-coding RNAs and coding RNAs. Despite the good performances reported in many studies for those tools, they generate a significant number of false positives and false negatives (7), and their reliability is questionable, which introduces uncertainty to the findings. As reported in the supplementary data (Figure S2), the lncRNA content of the LncPlankton database varies according to the coding potential prediction tool used; for instances only 7.73% of the transcripts of the MMETSP datasets were predicted as lncRNAs when using mRNN but this number reached 33.64% with LGC. Such difference in findings is inevitably caused by the intrinsic algorithms used by each method, the training set, and the fine-tuned learning features and parameters (see supplementary Table S1). In our pipeline, we opted for the use of a combination of multiple tools in a majority voting setting, inspired from the ensemble method technique used in machine learning. With this approach, 19% of the whole planktonic transcriptomes surveyed in this study was labelled as “lncRNAs”.

The choice of the coding potential prediction tool is not trivial and in most cases the criteria of evaluating these tools was based on the accuracy, sensitivity, specificity, and the computational time, but nothing has been reported about the reliability. In our case, the reliability of a predicted lncRNA transcript was calculated as the number of tools labelling the transcript as lncRNAs and dividing by 10, the total number of used tools. Therefore, a non-coding score value closer to 1 denotes a higher reliability. The density of this score, calculated on the lncRNAs found by our pipeline and distributed across groups shows high density at 0.6 and 0.7, decreasing at higher score values (Supplementary Figure S3). A significant part of lncRNAs found have a **medium** reliability of 0.6 or 0.7, meaning that at least 6 or 7 tools have predicted them as lncRNAs. A less significant number of lncRNAs have a score = 1 (239,116, ∼10,8% of the predicted lncRNAs), which means all the tools agreed that those transcripts are indeed lncRNAs. We called these **high confidence** lncRNAs. The remaining lncRNAs found have a **good** reliability with a score ranging between 0.8 and 0.9.

Regarding the filters introduced in our pipeline to discriminate lncRNA from other transcripts, default values were used for the transcript length (= 200nt), and the putative peptide length (= 100aa). Regarding peptide length, 1,475,973 lncRNA-like having predicted peptides > 3aa long which represent ∼ 67% of the total identified lncRNAs. Since lncRNAs are likely to possess ORFs purely by chance, discarding any transcripts containing ORFs would result in losing a significant number of true lncRNAs. Although this value has no fixed limit, the user can customize it in the SHINY prediction application developed and integrated to the LncPlankton user interface.

As part of our pipeline, we used a meta-learner combining multiple and diverse protein coding potential tools. We assessed the performance of our meta-learner using gold standard metrics such as the accuracy and AUC-ROC, and compared to each single tool. Overall, our method showed similar performance to the top performing coding potential tools, and it maintained a good performance in challenging cases in which some tools found difficulties to discriminate coding from non-coding transcripts. For highly imbalanced datasets like *B_taurus, X_tropicalis*, and *M_mulatta* which have a 1:72, 1:32, and 1:22 ratio of the number of coding to non-coding instances, some tools performed poorly longdist and RNAmining, while our method preserved a high accuracy meaning that our method is not impacted by the imbalancedness between coding and non-coding classes. The following aspects supported our choice to use the meta-learner proposed: **i)** the meta-learner used diverse and heterogeneous models covering a broad range of species since each model related to each tool was trained using transcripts of different species. Therefore, we believe that our method is a suitable choice for cross-species prediction; **ii)** our method achieved the highest accuracy. There are two reasons why the meta-learner outperforms others. First, our method benefited from the ensemble method principle since it combined multiple single tools. Second, the method integrated information from different types of features to enhance lncRNA prediction performances; **iii)** our method is robust showing a low variability of the prediction across the different testing datasets; **iv)** our method promotes the diversity which is coming from the different mathematics and algorithms used by each single tool, and fine-tuned features and parameters. Indeed, by pooling all the tools together, our method allowed an in-depth analysis of the transcripts by using all possible features describing the sequences; **v)** the reliability of the prediction in our method can be measured by the non-coding potential probability of the ensemble. It reflects the agreement of the tools to recognize the correct class of a transcript. In our method, we set the threshold to a value = 0.6 i.e. if at least 6 out of 10 tools recognized a transcript as non-coding RNA, it is more likely to be a non-coding RNA than a coding RNA. Increasing this threshold to 1 means that all the tools recognize it as a non-coding RNA. We believe the majority voting tool offers us a reliable choice for lncRNA identification, beyond what a single tool can offer at the moment.

We collected all the reliable lncRNAs predicted by our pipeline and their information, and stored them in a MySQL database termed LncPlankton. Many databases of lncRNAs have been implemented in the past few years but most of them are dedicated to humans, vertebrates and plants. To date, no other database of this scale collecting non-coding transcripts on planktonic organisms, with representatives from the major branches of the eukaryotic tree of life, has been developed. Although a few algal species have been included in plant lncRNA databases: 6 algae in GreenC (8), and 3 algae in Cantatadb (7), no database was completely dedicated to lncRNA in planktonic species. Compared with other lncRNA databases, the lncRNA transcripts of LncPlankton were predicted using the majority voting pipeline with stringent filters making them more reliable than what we could get with other pipelines using a single coding potential prediction tool.

Similar to other lncRNA databases, LncPlankton v1.0 includes all the basic information of a given lncRNA including the nucleotide sequence, the longest ORF sequence, and the predicted secondary structure. Two missing features of lncRNA recorded in LncPlankton v1.0 are the genomic coordinates (the start and stop position, and the strand information), and the expression profiles. This is due to the lack of an annotated genome for the species surveyed (11). In addition to the regular update of the database, the following future developments will be considered: **i)** the genomic coordinates of lncRNAs will be incorporated to further classify the lncRNAs identified as long intergenic ncRNAs (lincRNAs), intronic-lncRNAs, or antisense as this information becomes available; **ii)** the expression patterns can be integrated for the functional study of lncRNAs identified; **iii)** even if lncRNAs do not show high conservation at the nucleotide level, a sequence clustering using *Orthofinder* algorithm will be performed to provide information about highly conserved lncRNAs; **iv)** given that lncRNAs of related functions often share similar short motif profiles, we will use a kmer-based approach called *SEEKR* proposed in (34) to quantify the nonlinear sequence homology and the evolutionary relationships between the lncRNAs predicted in LncPlankton.

In addition, the UI will be improved accordingly by adding a submit page to the LncPlankton interface to enhance the interaction between the research community and the database, and to encourage all scientists to supply their own assembled contig data. All submitted entries will be processed by our procedure described in Materials and Methods.

## AVAILABILITY

The package of the majority voting procedure was implemented in R, and can be downloaded on LncPlankton website at https://www.lncplankton.bio.ens.psl.eu/files/tools/majorityLNC.tar.gz.The source code can be found at https://gitlab.com/a5076/majorityLNC. The database is freely available without restrictions for use by academics and non-commercial researchers. The web server is publicly available at https://www.lncplankton.bio.ens.psl.eu/. Inquiries concerning the database may be directed to.

## SUPPLEMENTARY DATA

Supplementary Data are available at bioRxiv online.

## ACKNOWLEDGEMENTS

This work has used the computational resources of the Bioclust cluster (https://bioclustg01.bioclust.biologie.ens.fr/). Hosting and logistic resources have been provided by the computational platform of the ENS Biology department.

