## Supplementary material for "LncPlankton V1.0: a comprehensive collection of plankton long non-coding RNAs"

**Supplementary Table S1**. Summary of coding potential computational approaches included in the proposed majority voting procedure

| Tool | Algorithm type | Training species | Classifying features | Can handle partial length transcripts? | Input file | Programming language | Ref. |
|---|---|---|---|---|---|---|---|
| **CPC2** | SVM | Human, mouse, fly and zebrafish | Fickett score, ORF length, ORF integrity, isoelectric point pI | NO | FASTA | Python3.5 | (1) |
| **LGC** | Maximum Likelihood Estimation | Human | ORF length, GC content | NO | FASTA/BED/ GTF | Python3.5 | (2) |
| **LncFinder** | SVM | Human, mouse and wheat | Length, ORF coverage, distance of hexamer ORF, secondary structure, physiochemical | NO | FASTA | R4.1.2 | (3) |
| **CPAT** | Logistic regression | human, mouse, fly and zebrafish | ORF size, ORF coverage, Fickett score, Hexamer score | NO | BED/FASTA | Python3.5 | (4) |
| **LncADeep** | Deep Learning (DBN) | Human and mouse | ORF length, ORF coverage, Entropy Density Profile (EDP) of ORFs, Hexamer score, UTR coverage, GC content, Fickett score, HMMER index, longest CDS | YES | FASTA | Python2.7<br><br>Backend engine = “theano” | (5) |
| **LncDC** | XGboost | Human and mouse | Sequence intrinsic features: GC content, ORF length, fickett score, etc. Secondary structure related features: Minimum free energy, SASS k-mer scores, etc. Protein features: pH isoelectric point, molecular weight, aromaticity, and stability index. | NO | FASTA | Python3.9 | (6) |
| **Longdist** | SVM | Human, mouse, and zebrafish | ORF length, relative length occurrences of k-mers | NO | FASTA | Python3.5 | (7) |
| **mRNN** | Deep Learning (RNN) | Human and mouse | k-mers | NO | FASTA | Python2.7 | (8) |
| **RNAmining** | XGboost | 15 species of | All combinations of trinucleotides | NO | FASTA | Python3.8 | (9) |

| | | | | | | | |
|---|---|---|---|---|---|---|---|
| | | distinct representative chordata clades including human and mouse | count: 64 features | | | | |
| **RNASamba** | Deep Learning (CNN/IGLOO) | Human | Max ORF length, k-mers, numeric representation of sequences | YES | FASTA | Python3.5/Rust<br><br>Backend engine = “tensorflow” | (10) |

**Supplementary Table S2**. The independent testing sets used for the evaluation of the coding potential tools

| Species | Data type | Number of transcripts | Source |
|---|---|---|---|
| **A_thaliana** | ncRNA | 5654 | https://rnamining.integrativebioinformatics.me |
| | Protein coding | 5654 | |
| **C_elegans** | ncRNA | 25279 | https://rnamining.integrativebioinformatics.me |
| | Protein coding | 25279 | |
| **C_auratus** | ncRNA | 7502 | https://rnamining.integrativebioinformatics.me |
| | Protein coding | 7502 | |
| **P_textilis** | ncRNA | 743 | https://rnamining.integrativebioinformatics.me |
| | Protein coding | 743 | |
| **R_norvegicus** | ncRNA | 9331 | https://rnamining.integrativebioinformatics.me |
| | Protein coding | 9331 | |
| **T_carolina** | ncRNA | 1027 | https://rnamining.integrativebioinformatics.me |
| | Protein coding | 1027 | |
| **G_gorilla** | ncRNA | 7989 | https://rnamining.integrativebioinformatics.me |
| | Protein coding | 7989 | |
| **D_melanogaster** | ncRNA | 3976 | https://rnamining.integrativebioinformatics.me |
| | Protein coding | 3976 | |
| **S_cerevisiae** | ncRNA | 413 | Ensembl |
| | Protein coding | 6713 | NCBI RefSeq |
| **D_rerio** | ncRNA | 10662 | Ensembl |
| | Protein coding | 15594 | NCBI RefSeq |
| **H_sapiens** | ncRNA | 22389 | Gencode |
| | Protein coding | 22389 | NCBI RefSeq |
| **M_musculus** | ncRNA | 6015 | Ensembl |
| | Protein coding | 6015 | NCBI RefSeq |
| **B_taurus** | ncRNA | 182 | Ensembl |
| | Protein coding | 13190 | NCBI RefSeq |
| **X_tropicalis** | ncRNA | 279 | Ensembl |
| | Protein coding | 8874 | NCBI RefSeq |
| **P_troglodytes** | ncRNA | 1166 | Ensembl |
| | Protein coding | 1906 | NCBI RefSeq |
| **M_mulatta** | ncRNA | 359 | Ensembl |
| | Protein coding | 5709 | NCBI RefSeq |
| **S_scrofa** | ncRNA | 241 | Ensembl |
| | Protein coding | 3978 | NCBI RefSeq |
| **P_abelii** | ncRNA | 392 | Ensembl |
| | Protein coding | 3401 | NCBI RefSeq |

**Supplementary Table S3**. Accuracy of the tools and the majority voting procedure evaluated on the independent testing sets (described in Table S2)

| | CPC2 | CPAT | LGC | LncADeep | LncDC | lncFinder | longdist | mRNN | RNAmining | RNASamba | Majority |
|---|---|---|---|---|---|---|---|---|---|---|---|
| **A_thaliana** | **0.92** | 0.90 | **0.92** | 0.80 | **0.92** | 0.88 | 0.89 | 0.84 | 0.90 | 0.82 | **0.92** |
| **B_taurus** | 0.96 | 0.98 | 0.93 | **0.99** | 0.98 | 0.97 | 0.32 | 0.98 | 0.26 | **0.99** | 0.98 |
| **C_elegans** | 0.96 | 0.94 | 0.95 | 0.75 | 0.97 | 0.87 | 0.98 | 0.96 | 0.63 | 0.96 | **0.99** |
| **C_auratus** | 0.86 | 0.87 | 0.85 | 0.78 | 0.83 | 0.85 | **0.88** | 0.86 | **0.88** | 0.84 | 0.84 |
| **D_melanogaster** | 0.92 | 0.92 | 0.90 | 0.85 | **0.93** | 0.90 | 0.89 | 0.81 | **0.93** | 0.76 | 0.92 |
| **G_gorilla** | 0.95 | 0.95 | 0.93 | 0.88 | 0.93 | 0.93 | 0.95 | 0.96 | 0.92 | **0.97** | **0.97** |
| **H_sapiens** | 0.93 | 0.94 | 0.91 | 0.94 | **0.97** | 0.93 | 0.56 | 0.92 | 0.54 | 0.93 | 0.95 |
| **M_mulatta** | 0.95 | 0.96 | 0.91 | **0.99** | 0.97 | 0.95 | 0.39 | 0.98 | 0.27 | **0.99** | 0.98 |
| **M_musculus** | 0.92 | 0.93 | 0.90 | 0.92 | 0.95 | 0.93 | 0.59 | 0.91 | 0.56 | 0.90 | **0.94** |
| **P_troglodytes** | 0.97 | 0.96 | 0.94 | 0.95 | **0.98** | 0.95 | 0.62 | 0.96 | 0.56 | **0.98** | **0.98** |
| **P_abelii** | 0.95 | 0.97 | 0.94 | **0.98** | **0.98** | 0.96 | 0.22 | **0.98** | 0.18 | **0.98** | 0.97 |
| **P_textilis** | **0.92** | 0.90 | 0.91 | 0.83 | 0.88 | 0.89 | **0.92** | 0.89 | **0.92** | 0.90 | 0.91 |
| **R_norvegicus** | 0.92 | 0.93 | 0.89 | 0.88 | 0.88 | 0.91 | 0.93 | 0.93 | 0.89 | 0.93 | **0.94** |
| **S_cerevisiae** | 0.89 | 0.83 | 0.90 | 0.92 | 0.89 | 0.81 | 0.99 | 0.94 | **0.97** | 0.95 | **0.97** |
| **S_scrofa** | 0.96 | 0.97 | 0.92 | **0.99** | 0.98 | 0.96 | 0.48 | 0.98 | 0.37 | **0.99** | 0.98 |
| **T_carolina** | 0.93 | 0.93 | 0.93 | 0.85 | 0.90 | 0.92 | 0.88 | **0.94** | 0.89 | **0.94** | 0.93 |
| **X_tropicalis** | 0.97 | 0.97 | 0.95 | **0.99** | 0.98 | 0.97 | 0.21 | 0.97 | 0.13 | **0.99** | 0.98 |
| **D_rerio** | **0.93** | **0.93** | 0.92 | 0.88 | **0.93** | **0.93** | 0.48 | 0.89 | 0.46 | 0.89 | 0.92 |

**Supplementary Table S4**. Sensitivity of the tools and the majority voting procedure evaluated on the independent testing sets (described in Table S2)

| | CPC2 | CPAT | LGC | LncADeep | LncDC | lncFinder | longdist | mRNN | RNAmining | RNASamba | Majority |
|---|---|---|---|---|---|---|---|---|---|---|---|
| **A_thaliana** | 0.93 | 0.90 | 0.93 | 0.98 | 0.95 | 0.87 | **1.00** | 0.97 | **1.00** | 0.98 | 0.99 |
| **B_taurus** | 0.96 | 0.98 | 0.93 | **0.99** | 0.98 | 0.97 | 0.31 | 0.98 | 0.25 | **0.99** | 0.98 |
| **C_elegans** | 0.92 | 0.89 | 0.90 | 0.98 | 0.97 | 0.85 | **1.00** | 0.98 | **1.00** | 0.98 | 0.99 |
| **C_auratus** | 0.97 | 0.97 | 0.96 | **1.00** | 0.97 | 0.95 | 0.93 | 0.99 | **1.00** | 0.99 | 0.99 |
| **D_melanogaster** | 0.95 | 0.97 | 0.94 | 0.99 | 0.99 | 0.93 | **1.00** | 0.99 | **1.00** | 0.99 | **1.00** |
| **G_gorilla** | 0.90 | 0.91 | 0.88 | 0.97 | 0.93 | 0.86 | 0.95 | 0.96 | **1.00** | 0.96 | 0.97 |
| **H_sapiens** | 0.95 | 0.96 | 0.93 | 0.98 | 0.98 | 0.96 | 0.16 | 0.97 | 0.12 | **0.99** | 0.97 |
| **M_mulatta** | 0.95 | 0.96 | 0.91 | **1.00** | 0.97 | 0.95 | 0.36 | 0.98 | 0.23 | 0.99 | 0.98 |
| **M_musculus** | 0.96 | 0.97 | 0.94 | **0.99** | 0.98 | 0.97 | 0.21 | 0.97 | 0.16 | **0.99** | 0.98 |
| **P_troglodytes** | 0.95 | 0.94 | 0.91 | **0.98** | 0.96 | 0.93 | 0.44 | 0.95 | 0.35 | **0.98** | 0.96 |
| **P_abelii** | 0.95 | 0.97 | 0.93 | 0.98 | 0.98 | 0.96 | 0.15 | 0.98 | 0.10 | **0.99** | 0.97 |
| **P_textilis** | 0.93 | 0.91 | 0.91 | **1.00** | 0.93 | 0.89 | 0.94 | 0.98 | **1.00** | 0.98 | 0.98 |
| **R_norvegicus** | 0.90 | 0.94 | 0.85 | 0.99 | 0.88 | 0.90 | 0.96 | 0.98 | **1.00** | 0.97 | 0.98 |
| **S_cerevisiae** | 0.89 | 0.82 | 0.90 | 0.95 | 0.89 | 0.79 | **0.99** | 0.94 | 0.98 | 0.95 | 0.97 |
| **S_scrofa** | 0.95 | 0.97 | 0.91 | **1.00** | 0.98 | 0.96 | 0.46 | 0.98 | 0.35 | 0.99 | 0.98 |
| **T_carolina** | 0.91 | 0.92 | 0.89 | 0.99 | 0.91 | 0.89 | 0.90 | 0.97 | **1.00** | 0.96 | 0.97 |
| **X_tropicalis** | 0.97 | 0.97 | 0.94 | **1.00** | 0.98 | 0.97 | 0.20 | 0.97 | 0.12 | **1.00** | 0.98 |
| **D_rerio** | 0.97 | 0.97 | 0.94 | **1.00** | 0.99 | 0.97 | 0.31 | 0.98 | 0.18 | 0.99 | 0.98 |

**Supplementary Table S5**. Specificity of the tools and the majority voting procedure evaluated on the independent testing sets (described in Table S2)

| | CPC2 | CPAT | LGC | LncADeep | LncDC | lncFinder | longdist | mRNN | RNAmining | RNASamba | Majority |
|---|---|---|---|---|---|---|---|---|---|---|---|
| **A_thaliana** | **0.91** | 0.90 | **0.91** | 0.62 | 0.89 | 0.90 | 0.75 | 0.70 | 0.81 | 0.66 | 0.84 |
| **B_taurus** | **1.00** | 0.98 | **1.00** | 0.82 | **1.00** | 0.99 | 0.82 | 0.95 | 0.82 | 0.86 | 0.99 |
| **C_elegans** | **1.00** | 0.98 | **1.00** | 0.52 | 0.98 | 0.89 | 0.79 | 0.94 | 0.27 | 0.95 | 0.96 |
| **C_auratus** | 0.76 | 0.77 | 0.75 | 0.55 | 0.57 | 0.76 | **0.79** | 0.72 | 0.75 | 0.69 | 0.55 |
| **D_melanogaster** | **0.88** | 0.87 | 0.87 | 0.71 | 0.84 | 0.87 | 0.75 | 0.62 | 0.85 | 0.53 | 0.81 |
| **G_gorilla** | **0.99** | 0.98 | **0.99** | 0.78 | 0.97 | **0.99** | 0.92 | 0.96 | 0.84 | 0.98 | 0.97 |
| **H_sapiens** | 0.91 | 0.91 | 0.88 | 0.89 | 0.96 | 0.91 | **0.96** | 0.88 | **0.96** | 0.87 | 0.94 |
| **M_mulatta** | **1.00** | 0.99 | **1.00** | 0.92 | **1.00** | 0.99 | 0.90 | 0.97 | 0.87 | 0.94 | **1.00** |
| **M_musculus** | 0.89 | 0.88 | 0.87 | 0.86 | 0.93 | 0.89 | 0.96 | 0.85 | **0.95** | 0.80 | 0.91 |
| **P_troglodytes** | **1.00** | 0.99 | **1.00** | 0.90 | **1.00** | **1.00** | 0.93 | 0.97 | 0.91 | 0.97 | **1.00** |
| **P_abelii** | **1.00** | 0.99 | **1.00** | 0.90 | **1.00** | 0.99 | 0.88 | 0.98 | 0.87 | 0.94 | **1.00** |
| **P_textilis** | **0.91** | 0.89 | **0.91** | 0.67 | 0.75 | 0.90 | 0.87 | 0.81 | 0.83 | 0.82 | 0.70 |
| **R_norvegicus** | **0.93** | **0.93** | **0.93** | 0.77 | 0.87 | **0.93** | 0.87 | 0.88 | 0.79 | 0.88 | 0.84 |
| **S_cerevisiae** | **1.00** | 0.97 | 0.99 | 0.36 | 0.96 | **1.00** | 0.83 | 0.89 | 0.74 | 0.96 | 0.98 |
| **S_scrofa** | 0.98 | 0.95 | 0.98 | 0.83 | **0.99** | 0.95 | 0.83 | 0.90 | 0.80 | 0.87 | 0.95 |
| **T_carolina** | **0.96** | 0.95 | **0.96** | 0.72 | 0.86 | 0.95 | 0.80 | 0.90 | 0.78 | 0.92 | 0.76 |
| **X_tropicalis** | **1.00** | **1.00** | **1.00** | 0.72 | **1.00** | **1.00** | 0.89 | 0.95 | 0.66 | 0.94 | **1.00** |
| **D_rerio** | **0.88** | 0.86 | 0.87 | 0.70 | 0.82 | 0.86 | 0.86 | 0.75 | **0.88** | 0.73 | 0.79 |

**Supplementary Table S6**. AUC performance of the tools and the majority voting procedure evaluated on the independent testing sets (described in Table S2)

| | CPC2 | CPAT | LGC | LncADeep | LncDC | lncFinder | longdist | mRNN | RNAmining | RNASamba | Majority |
|---|---|---|---|---|---|---|---|---|---|---|---|
| **A_thaliana** | 0.95 | 0.94 | 0.89 | 0.86 | 0.94 | 0.91 | 0.95 | 0.91 | 0.41 | 0.90 | **0.99** |
| **B_taurus** | **1.00** | **1.00** | 0.99 | 0.98 | **1.00** | **1.00** | 0.61 | 0.99 | 0.26 | 0.99 | **1.00** |
| **C_elegans** | 0.99 | 0.99 | 0.99 | 0.87 | **1.00** | 0.89 | 0.97 | 0.99 | 0.10 | 0.99 | **1.00** |
| **C_auratus** | 0.92 | **0.93** | 0.84 | 0.92 | 0.85 | 0.88 | 0.90 | 0.94 | 0.46 | 0.91 | 0.92 |
| **D_melanogaster** | 0.96 | 0.96 | 0.83 | 0.94 | 0.95 | 0.93 | 0.94 | 0.96 | 0.61 | 0.92 | **0.99** |
| **G_gorilla** | 0.99 | 0.99 | 0.98 | 0.95 | 0.99 | 0.95 | 0.97 | 0.98 | 0.33 | 0.99 | **1.00** |
| **H_sapiens** | 0.97 | 0.97 | 0.91 | 0.96 | **0.99** | 0.97 | 0.63 | 0.96 | 0.44 | 0.97 | 0.98 |
| **M_mulatta** | **1.00** | 0.99 | 0.98 | 0.99 | **1.00** | 0.99 | 0.71 | 0.99 | 0.26 | **1.00** | **1.00** |
| **M_musculus** | 0.97 | 0.96 | 0.86 | 0.96 | **0.98** | 0.96 | 0.69 | 0.95 | 0.51 | 0.95 | **0.98** |
| **P_troglodytes** | **1.00** | 0.99 | 0.99 | 0.98 | **1.00** | 0.99 | 0.81 | 0.98 | 0.31 | 0.99 | 0.99 |
| **P_abelii** | **1.00** | 0.99 | **1.00** | 0.99 | **1.00** | **1.00** | 0.53 | 0.99 | 0.15 | **1.00** | **1.00** |
| **P_textilis** | 0.95 | 0.95 | 0.91 | 0.92 | 0.89 | 0.91 | 0.95 | 0.95 | 0.35 | 0.95 | **0.96** |
| **R_norvegicus** | 0.95 | 0.96 | 0.89 | 0.95 | 0.93 | 0.94 | 0.95 | 0.97 | 0.39 | 0.96 | **0.98** |
| **S_cerevisiae** | 0.98 | 0.96 | 0.97 | 0.65 | 0.99 | 0.93 | 0.98 | 0.96 | 0.15 | 0.97 | **1.00** |
| **S_scrofa** | **1.00** | 0.98 | 0.96 | 0.96 | **1.00** | 0.98 | 0.76 | 0.97 | 0.35 | 0.96 | 0.99 |
| **T_carolina** | **0.97** | **0.97** | 0.95 | 0.93 | 0.92 | 0.93 | 0.89 | 0.96 | 0.31 | 0.96 | **0.97** |
| **X_tropicalis** | **1.00** | **1.00** | **1.00** | 0.96 | **1.00** | **1.00** | 0.62 | 0.98 | 0.10 | **1.00** | **1.00** |
| **D_rerio** | **0.96** | 0.95 | 0.91 | 0.93 | 0.94 | 0.95 | 0.70 | 0.93 | 0.33 | 0.92 | 0.94 |

**Supplementary Table S7**. Species in LncPlankton presenting the highest and lowest number of lncRNAs detected by group using the majority voting procedure

| Group | Species/Strain | Total transcripts | Total lncRNAs | High-confidence lncRNAs |
|---|---|---|---|---|
| Bacillariophyta | Pseudo_nitzschia_delicatissima_B596 | 19906 | 1009 | 21 |
| | Fragilariopsis_kerguelensis_L26_C5 | 75135 | 21011 | 1979 |
| Cercozoa | Minchinia_chitonis | 324 | 83 | 10 |
| | Lotharella_globosa_Strain_CCCM811 | 27107 | 5522 | 665 |
| Chlorophyta | Pycnococcus_sp_Strain_CCM1998 | 1060 | 201 | 7 |
| | Mantoniella_sp_Strain_CCMP1436 | 25653 | 9332 | 3190 |
| Ciliophora | Strombidium_rassoulzadegani_Strain_ras09 | 11186 | 1671 | 184 |
| | Condylostoma_magnum | 26175 | 20463 | 7719 |
| Cryptophyta | Hanusia_phi | 22358 | 1622 | 75 |
| | Geminigera_cryophila_Strain_CCMP2564 | 53848 | 15757 | 2280 |
| Dinophyta | Thoracosphaera_heimii_Strain_CCCM_670_CCMP_1069 | 92 | 10 | 0 |
| | Karenia_brevis_Wilson | 123091 | 23686 | 2883 |
| Haptophyta | Phaeocystis_cordata | 3701 | 642 | 31 |
| | Undescribed_sp_CCMP2436 | 48766 | 12923 | 3136 |
| Other-phyla | Mayorella_sp_Strain_BSH_02190019 | 1945 | 403 | 19 |
| | Eutreptiella_gymnastica_like_Strain_CCMP1594 | 74436 | 14439 | 3215 |
| Other-stramenopiles | Cafeteria_roenbergensis | 17122 | 778 | 9 |
| | Undescribed_sp_CCMP2098 | 85238 | 20282 | 1872 |

**Supplementary Figure S1**. A flowchart depicting the different steps of the pipeline used for the transcriptome assembly of the two species *Phaeodactylum tricornutum* and *Thalasiossira pseudonana.* The contigs < 150 bp were discarded to produce the final assembly fasta file for these two species as done in [11].

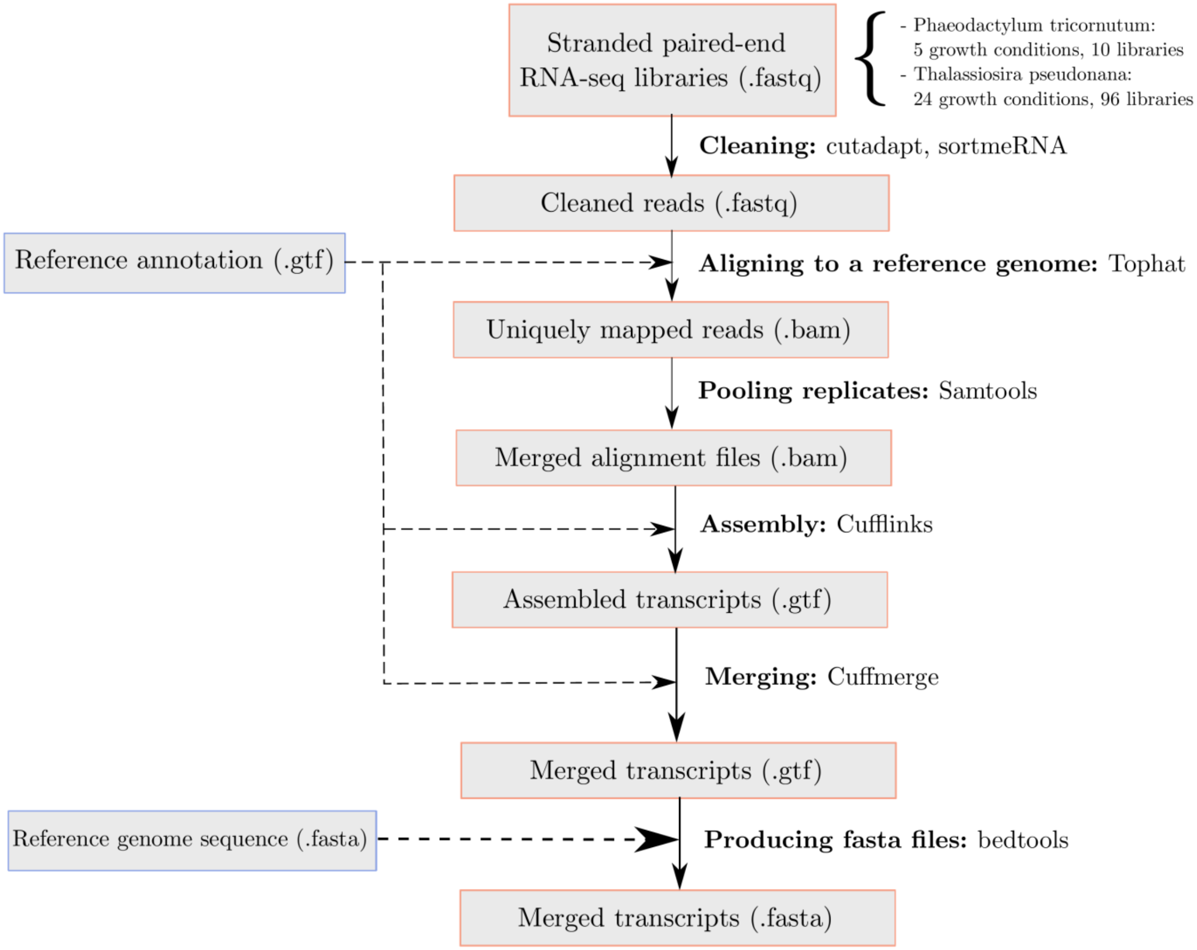

**Supplementary Figure S2**. The percentage of lncRNAs detected by each coding potential prediction method. The numbers corresponding to the percentages can be read on the y-axis.

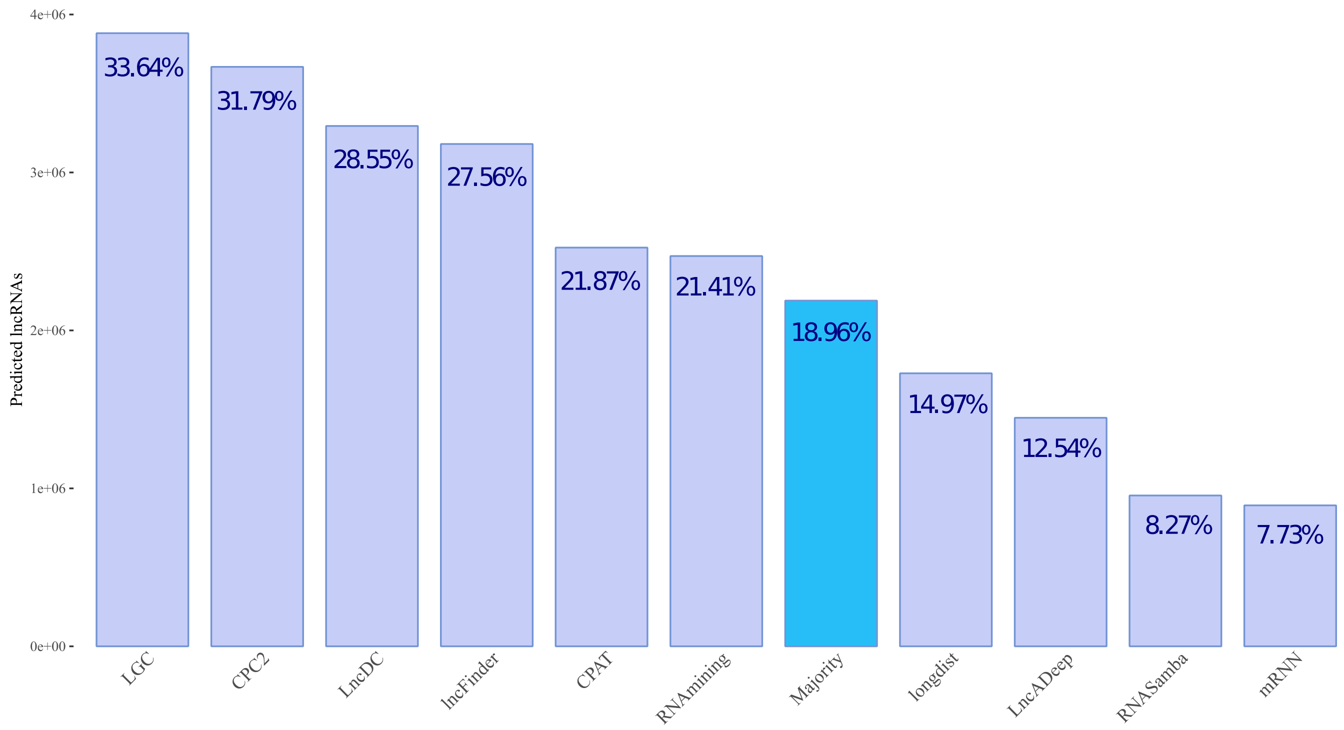

**Supplementary Figure S3**. Density of $(1 - coding\ potential\ probability)$ of lncRNAs in LncPlankton calculated by the majority voting procedure and distributed across the groups.

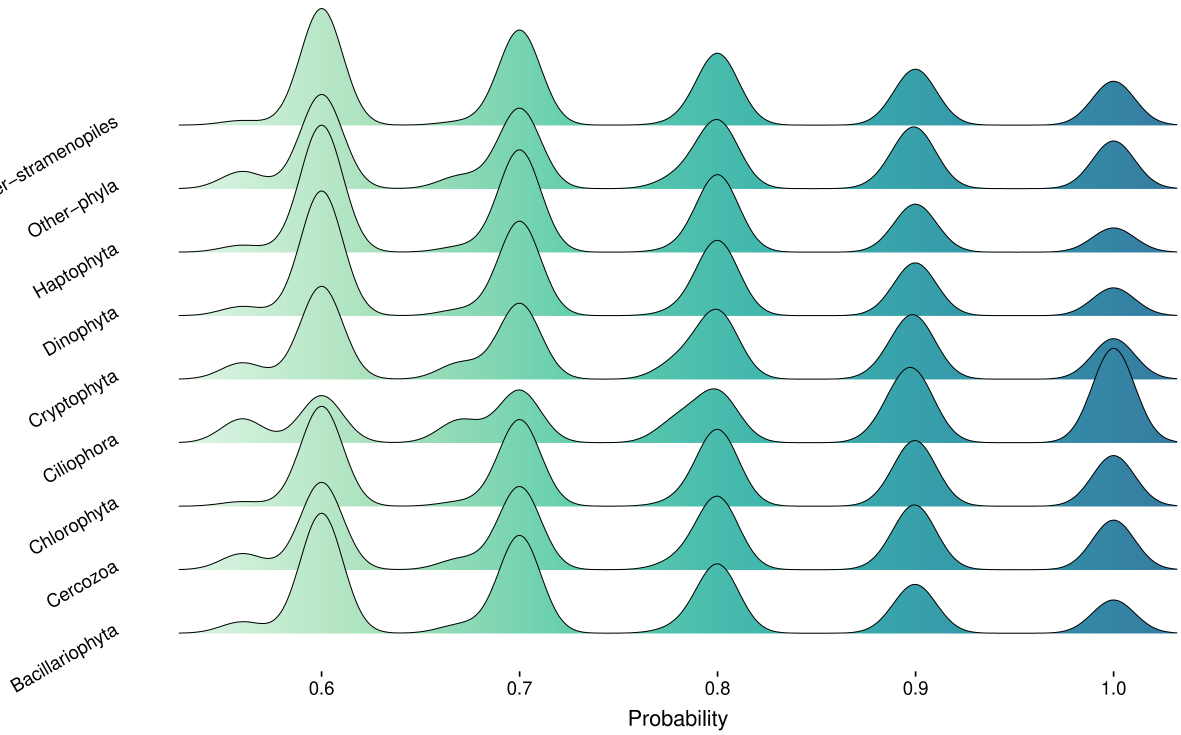
